# Identifying the internalization pathways of magnetotactic bacteria and magnetosomes by cancer cells

**DOI:** 10.1101/2025.11.26.690460

**Authors:** Lucía Gandarias, Lide Arana, Alicia G. Gubieda, Ana J. Pérez-Berná, Ana Abad-Díaz-de-Cerio, M. Luisa Fdez-Gubieda, Alicia Muela, Ana García-Prieto

## Abstract

Given the need for new tumor treatment strategies, therapies using magnetic nanoparticles and bacteria are gaining momentum. In this context, magnetotactic bacteria and magnetosomes could act as theranostic agents for use in magnetic hyperthermia, targeted drug delivery, and magnetic resonance imaging. Thus, understanding their interaction with target cells is essential to ensure their theranostic efficiency. This study investigates the uptake of magnetotactic bacteria (MSR-1) and magnetosomes by lung carcinoma cells (A549). First, MSR-1 and magnetosomes are imaged inside A549 cells using cryo soft X-ray tomography, revealing the presence of MSR-1 and magnetosomes inside endosomes. Subsequently, the endocytosis pathways involved in the internalization of MSR-1 and magnetosomes by the cells are elucidated. It is observed that MSR-1 mainly enter cells by receptor-mediated endocytosis, as described previously for other intracellular bacteria. However, the endocytosis of magnetosomes occurs mainly via phagocytosis or macropinocytosis, probably due to the large size of the formed magnetosome clusters. These findings fill a key gap in our understanding of the internalization of MSR-1 bacteria and magnetosomes by lung carcinoma cells, and establish a method applicable to studying their internalization by other target cell types.

**Graphical Abstract:** Magnetotactic bacteria (*Magnetospirillum gryphiswaldense* MSR-1) and magnetosomes, which are proposed as cancer theranostic mediators, are imaged inside A549 cancer cells in 3D using cryo soft X-ray tomography. Moreover, the endocytosis pathways that the cells use to internalize MSR-1 and magnetosomes are elucidated. MSR-1 predominantly enter A549 cells via receptor-mediated endocytosis, while magnetosomes are preferentially internalized via receptor-independent endocytosis.

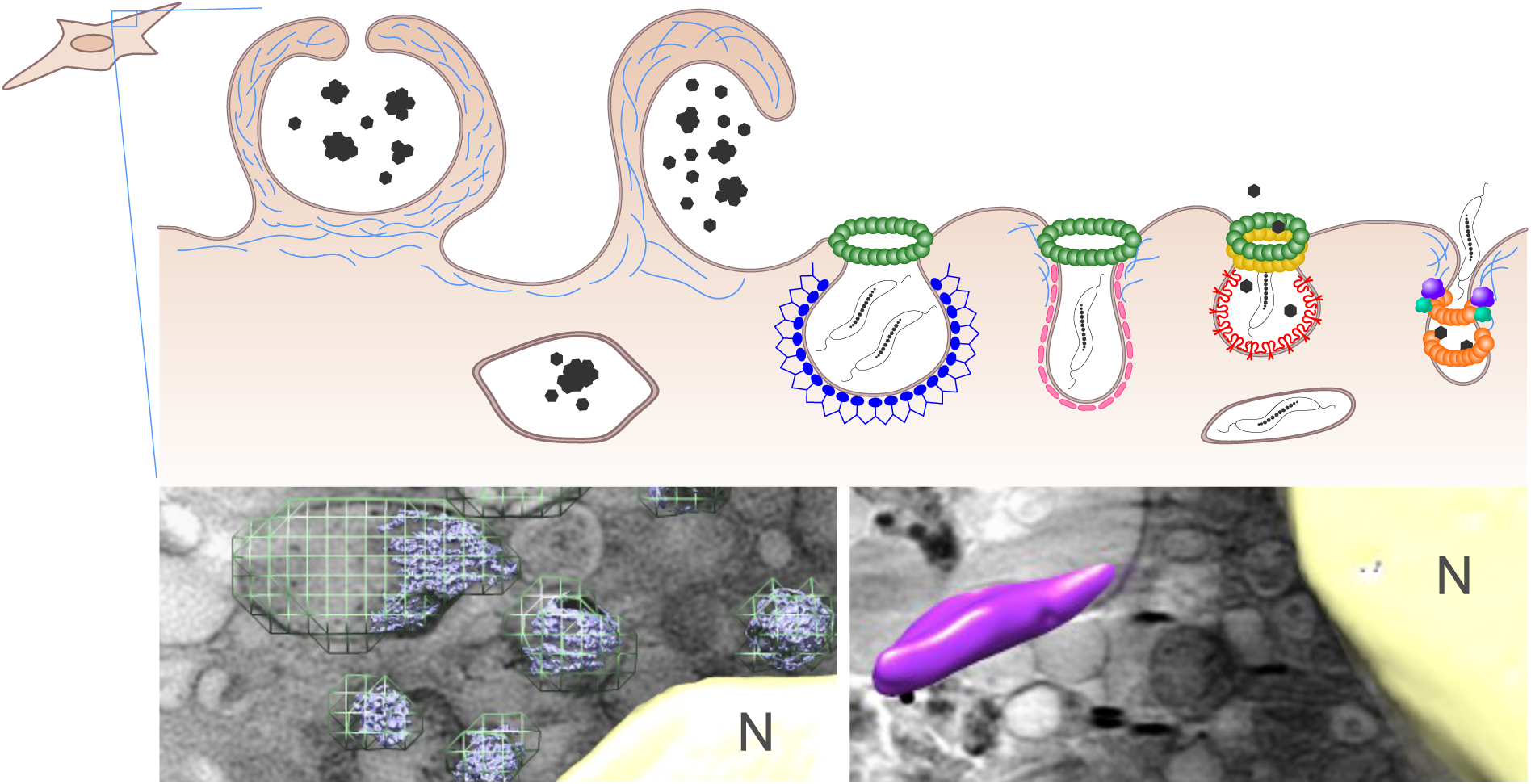

## Introduction

Challenges with current anti-tumour therapies, such as adverse side effects, difficulty in treating deep tumours, and induced tumour resistance to certain drugs, have led to the emergence of alternative approaches to treat cancer. One of the most innovative yet early-proposed strategies involves the use of bacteria. The first use of these microorganisms to treat cancer dates from the 19^th^ century when tumour regression in patients treated with *Streptococcus pyogenes* and *Serratia marcescens* was reported ^1,2^. However, with the emergence of other treatments such as chemotherapy or radiotherapy, bacterial-mediated therapy was left aside. From the current perspective of going towards a more specific cancer treatment to reduce side effects, bacteria are being revisited as potential therapy mediating systems. Currently, the use of an attenuated strain of *Mycobacterium tuberculosis* is used to treat patients with superficial, non-muscle invasive bladder cancer in what is known as the Bacillus Calmette-Guérin (BCG) treatment ^3^. Indeed, certain bacteria are immunostimulatory and can accumulate and/or thrive preferably in tumour microenvironments due to the hypoxic conditions and the lower immune surveillance as compared to other tissues. Moreover, bacteria can be used to deliver payloads such as drugs, immunomodulators, cytokines, or small interfering RNAs (siRNAs) to the tumour microenvironment ^4^.

One of the main challenges of using bacteria for cancer treatment is the ability to guide, actuate, and track them inside the body. For instance, bacteria can be genetically engineered to synthesize gas vesicles ^5^, or can be made magnetic using synthetic biology techniques ^6–8^ or by decorating them with magnetic nanoparticles ^9,10^, in order to be guided, actuated, and detected by focused ultrasound or magnetic fields, respectively.

Magnetotactic bacteria (MTB) are environmental Gram negative aquatic microorganisms that have the particularity of being able to synthesize magnetic nanoparticles named magnetosomes that they arrange into one or more chains ^11,12^. Due to their non-pathogenicity, motility, and magnetic properties, they have been proposed as theranostic microrobots ^13^ that can be guided using external magnetic fields ^14–17^, actuated with alternating magnetic fields in magnetic hyperthermia therapy ^18–22^, and tracked with magnetic resonance imaging (MRI) ^23–25^ and magnetic particle imaging (MPI) ^26^.

However, before MTB were proposed as theranostic agents, most of the research was performed on the use of isolated magnetosomes as mediators in similar applications ^27–33^. If the performance of MTB and magnetosomes is compared, there are strengths and limitations of both systems: on the one hand, MTB are motile thanks to their flagella and have a lower agglomeration tendency when compared to magnetosomes, but on the other hand, magnetosome agglomeration is beneficial to be detected using MPI as demonstrated by Makela et al. ^26^. In terms of magnetic material accumulation, cancer cells are able to internalize more magnetite when in contact with magnetosomes than with magnetotactic bacteria. Therefore, the use of one or the other system should be thought through depending on the application.

Ultimately, the ability of target cells to internalize and/or interact with MTB or magnetosomes will determine the efficiency of the application. Wang et al. ^34^ already addressed the endocytosis of magnetosomes by hepatic cancer cells, but the internalization of MTB has yet to be explored. This study aims to fill this gap by studying the interaction of MTB and magnetosomes with cancer cells using bacteria and magnetosomes from the species *Magnetospirillum gryphiswaldense* MSR-1 and lung carcinoma cells from the A549 cell line as models. The interaction between MSR-1/magnetosomes and A549 cells was observed using cryo soft X-ray tomography, a synchrotron based technique that allows for the 3D imaging of intracellular structures without the need to perform ultrathin sectioning or staining. Finally, the endocytosis pathways used by A549 cells to internalize MSR-1/magnetosomes were elucidated by using specific pharmacological inhibitors of endocytosis, combined with flow cytometry.

## Results and discussion

### Imaging MSR-1/magnetosome-cell interactions by cryo soft X-ray tomography

In a previous publication we observed MSR-1 bacteria (Figure S1A) inside A549 cells by using fluorescence and transmission electron microscopy (TEM) ^20^. The internalization of magnetosomes (Figure S1B) by several cell lines including cancer cells, macrophages, and mesenchymal stem cells has already been shown by various groups including ours using the aforementioned microscopic techniques ^28,29,35–38^. Both bright-field/fluorescence microscopy and TEM are widely used techniques, as they are commonly available in research laboratories; however, they do present certain limitations. Bright-field/fluorescence microscopy is a convenient technique for a first observation of cells but the resolution is low and it only allows the intracellular structures to be visualized if they are labeled using a fluorescent marker, for instance. On the other hand, TEM has a higher resolution and enables the observation of the different organelles but it requires chemical fixation and ultrathin sectioning of the samples as eukaryotic cells are too thick for electrons to penetrate. To overcome these drawbacks, in this work, cryo soft X-ray tomography (cryo-SXT) is used to image MSR-1 and magnetosomes inside A549 cells. This synchrotron based technique offers many advantages compared to the aforementioned techniques, such as the fact that it enables the full volume of the cells to be imaged, as opposed to the ultrathin sections observed in TEM. Furthermore, the sample preparation is more straightforward as it only requires ice-vitrification to preserve cells in a near-native state and to measure them in cryo to reduce radiation damage during data collection ^39,40^. When imaged at the so-called *water window* (284 - 543 eV), which lies between the inner-shell absorption edges of carbon and oxygen, soft X-rays can penetrate water layers up to 10 µm thick, enabling high-contrast visualization of carbon-rich structures. Under these conditions, a spatial resolution of approximately 40 nm can be achieved ^41^. This technique has previously been used to observe metal nanoparticles inside cells ^42,43^, magnetosomes inside magnetotactic bacteria ^44,45^, and intracellular structures ^46,47^. Here, it is used to observe MSR-1 and magnetosomes inside cancer cells.

Briefly, to prepare the samples, A549 cells were seeded on well plates containing carbon-coated TEM grids and incubated overnight to let them attach. The next day, MSR-1 and magnetosomes were added suspended in cell culture media and co-incubated with the cells for 24 h (MSR-1) or 2 h (magnetosomes). Then, the grids were washed with PBS before being mounted vertically on the plunge freezer and fiducials (100 nm gold nanoparticles for tomography reconstruction) were added before vitrification. The samples were kept under cryogenic conditions before being measured at the MISTRAL beamline of the ALBA synchrotron. More details on sample preparation and image acquisition can be found on the Materials and Methods section.

Figure 1 shows X-ray images and z-slices obtained in the *water window* (520 eV) of A549 cells containing MSR-1 (A-E) or magnetosomes (F-I). Figures 1A and F show an X-ray image of the cell and the red box marks the selected region for tomogram collection in each case. The images in Figure 1B and G are z-slices of the reconstructed tomogram corresponding to the region marked in Figure 1A and F, respectively. In Figure 1B MSR-1 can be distinguished within the cell cytoplasm in some of the slices (marked by yellow arrows). As opposed to TEM imaging of ultrathin cellular sections, where MSR-1 were only inferred inside the cells thanks to the more electrodense magnetosomes ^20^, cryo-SXT allows for the observation of the full bacterial body because of the natural contrast provided by MSR-1 imaged at the *water window*. Magnetosome chains can also be observed inside MSR-1 as shown in Figure 1C (marked with a turquoise arrow). Figure 1D shows a representative z-slice where additional cell structures have been identified (nucleus, N, mitochondria, M, lipid droplets, LD, and cellular vesicles, V). In some cases, MSR-1 can be observed embedded in endosomes (marked with green arrows), suggesting endocytosis as the main internalization mechanism. A volume segmentation of the bacteria from the tomogram (Figure 1E) provides 3D views of MSR-1 inside the cell adopting different orientations.

**Figure 1:**
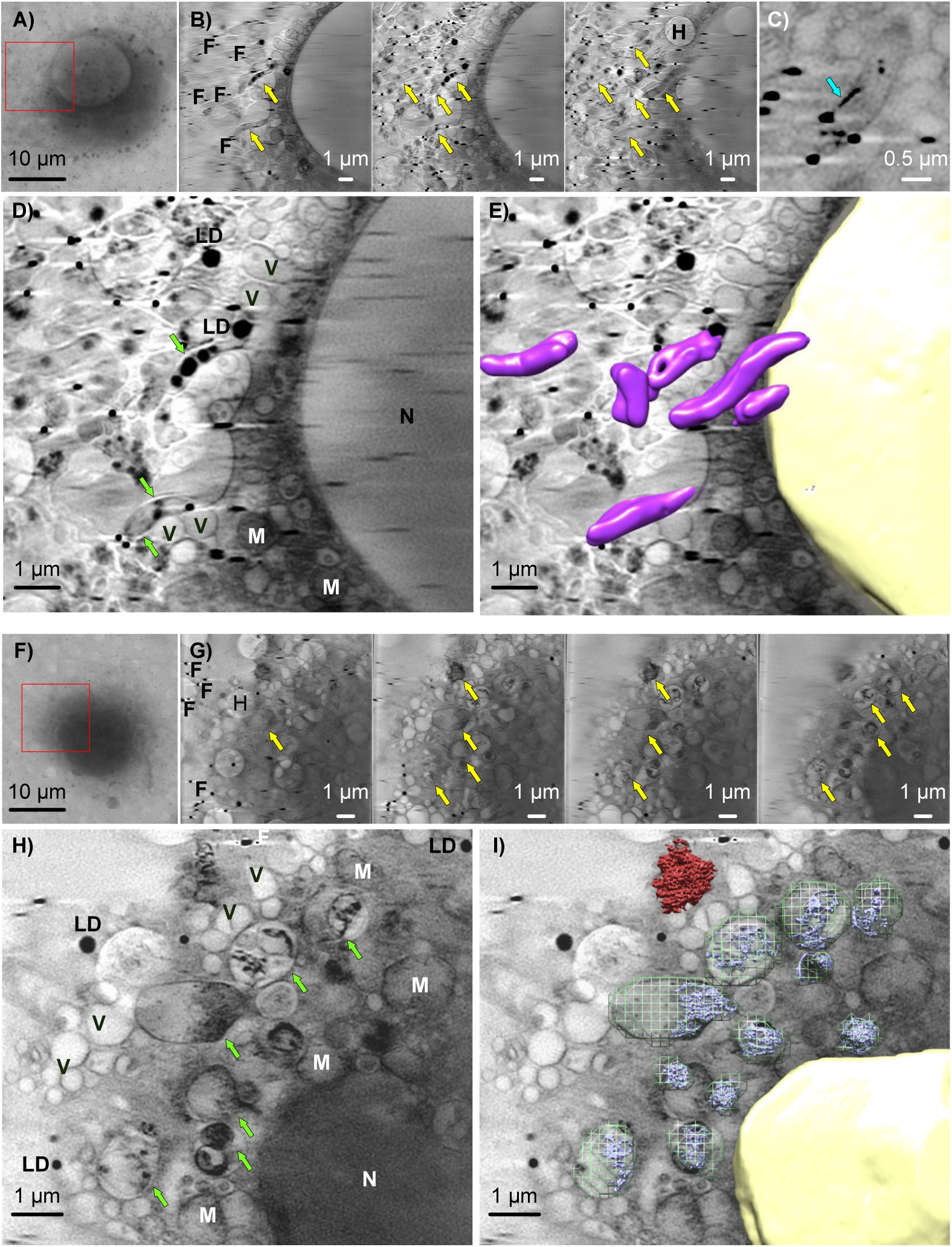
Cryo-SXT images of A549 cells loaded with MSR-1 (A-E) and magnetosomes (F-I). A,F) X-ray images of the measured cells, with the region chosen for tomogram collection highlighted by a red square. B,G) Z-slices of the reconstructed tomograms spaced 260 nm. The yellow arrows point at MSR-1 and magnetosome clusters. Carbon film holes are marked H and fiducials are marked F. C) Detail of a z-slice showing a MSR-1 bacterium inside the cell where the magnetosome chain is visible and marked with a turquoise arrow. D,E,H,I) Selected cryo-SXT tomogram slices (D,H) and segmented volumes representation (E,I) of the tomograms highlighting cell nuclei (yellow), MSR-1 (magenta, E), magnetosomes inside endosomes (blue, I), endosomes (green mesh, I), and magnetosomes attached to the plasma membrane (red, I). The observed intracellular structures are marked as: N, cell nuclei; M, mitochondria; LD, lipid droplets; V, vesicles; green arrows, endosomes containing MSR-1 or magnetosomes.

In the case of magnetosome-loaded cells (Figures 1F-I), magnetosomes form clusters that are clearly distinguished due to their high density (marked with yellow arrows in Figure 1G). Magnetosomes are observed within endosomes in the cell cytoplasm (marked with green arrows in Figure 1H), with variable sizes ranging between 0.7 and up to 3 µm. There is also a cluster that is attached to the cell membrane highlighted in red in Figure 1I. From the volume segmentation of the endosomes and the magnetosome clusters from the tomogram (Figure 1I), it can be observed that endosomes are only partially filled with magnetosomes, with a magnetosome-to-vesicle volume ratio up to 30%. The total magnetosome mass in the imaged volume can be estimated from the density of magnetite (5.24 g cm^-3^) as 11.7 pg, in accordance with the uptaken magnetite mass per cell estimated by magnetometry (64 pg ^36^), assuming a correction factor given the partial volume analyzed by cryo-SXT.

Further tomograms and videos of the 3D volume reconstructions can be found in the Supplementary Information section (Figure S2 and the Supplementary Videos).

### Elucidating the internalization pathways for MSR-1 and magnetosome uptake

Endocytosis is the primary mechanism for the internalization of nanoparticles and microorganisms into cells ^48,49^. According to the current classification there are six different endocytosis pathways that depend on the proteins involved in the process: phagocytosis, macropinocytosis, clathrin-mediated endocytosis, fast endophilin-mediated endocytosis (FEME), caveolae-dependent endocytosis, and clathrin-independent carrier glycosylphosphatidylinositol-anchored protein-enriched early endocytic compartment endocytosis (CLIC/GEEC) ^48,49^ (Figure 2). Independently of the pathway used for endocytosis, the cells first store the cargo in early endosomes that can be redirected to the cell membrane to excrete the content to the external milieu or that can fuse with lysosomes to proceed to the degradation of the cargo.

**Figure 2:**
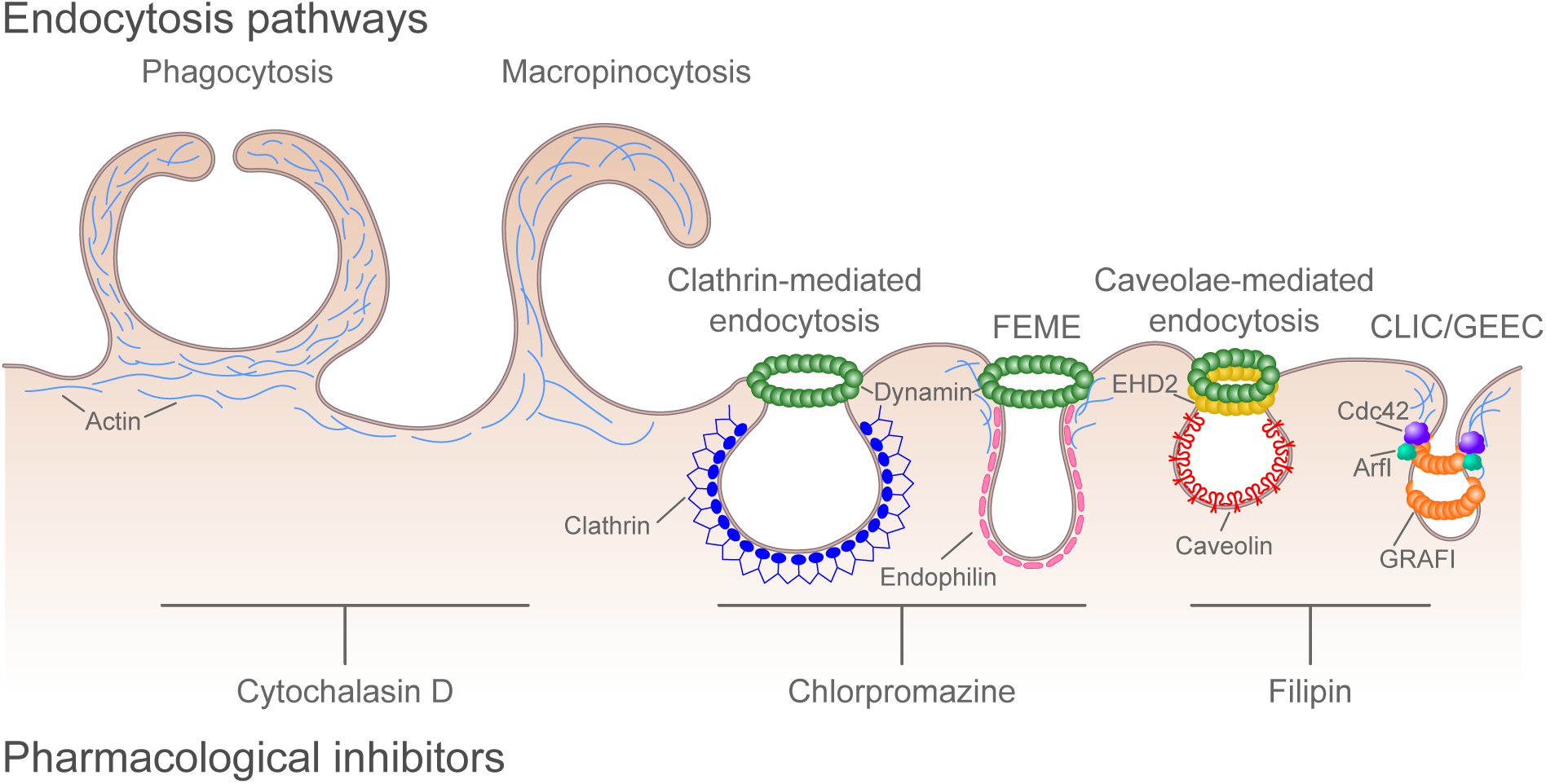
Overview of the endocytic pathways present in eukaryotic cells along with the specific pharmacological inhibitors used in the present work. FEME (fast endophilin-mediated endocytosis), CLIC/GEEC (clathrin-independent carrier glycosylphosphatidylinositol-anchored protein-enriched early endocytic compartment endocytosis), EHD2 (Eps15-homology domain-containing protein 2), Cdc42 (cell division cycle 42), ArfI (ADP-ribosylation factor 1), GRAFI (GTPase regulator associated with focal adhesion kinase-1).

To elucidate the particular endocytic route involved in the internalization of MSR-1/magnetosomes, three specific pharmacological inhibitors were tested (Figure 2). Cytochalasin D inhibits phagocytosis and macropinocytosis by interfering with actin polymerization necessary for both pathways ^50,51^. Chlorpromazine inhibits clathrin-mediated endocytosis preventing the assembly of clathrin-coated pits at the inner surface of the plasma membrane ^52^ and can also inhibit FEME as it prevents the formation of endophilin-positive assemblies needed for this pathway to take place ^53^. Filipin inhibits caveolae-dependent endocytosis by binding to sterols such as cholesterol and by disorganizing caveolin ^54^ but it has also been shown to inhibit the CLIC/GEEC pathway ^48^.

First, the inhibitor concentration that did not cause toxicity in the cells was established. For this, cell viability at increasing inhibitor concentrations was studied using flow cytometry and propidium iodide staining as detailed in the Materials and Methods section. The obtained results are represented in Figure 3 where the viability values are normalized to those of control cells (without inhibitor). In view of these results, to avoid any side effects, the inhibitor concentrations shown in the table of Figure 3 were used for the subsequent experiments.

**Figure 3:**
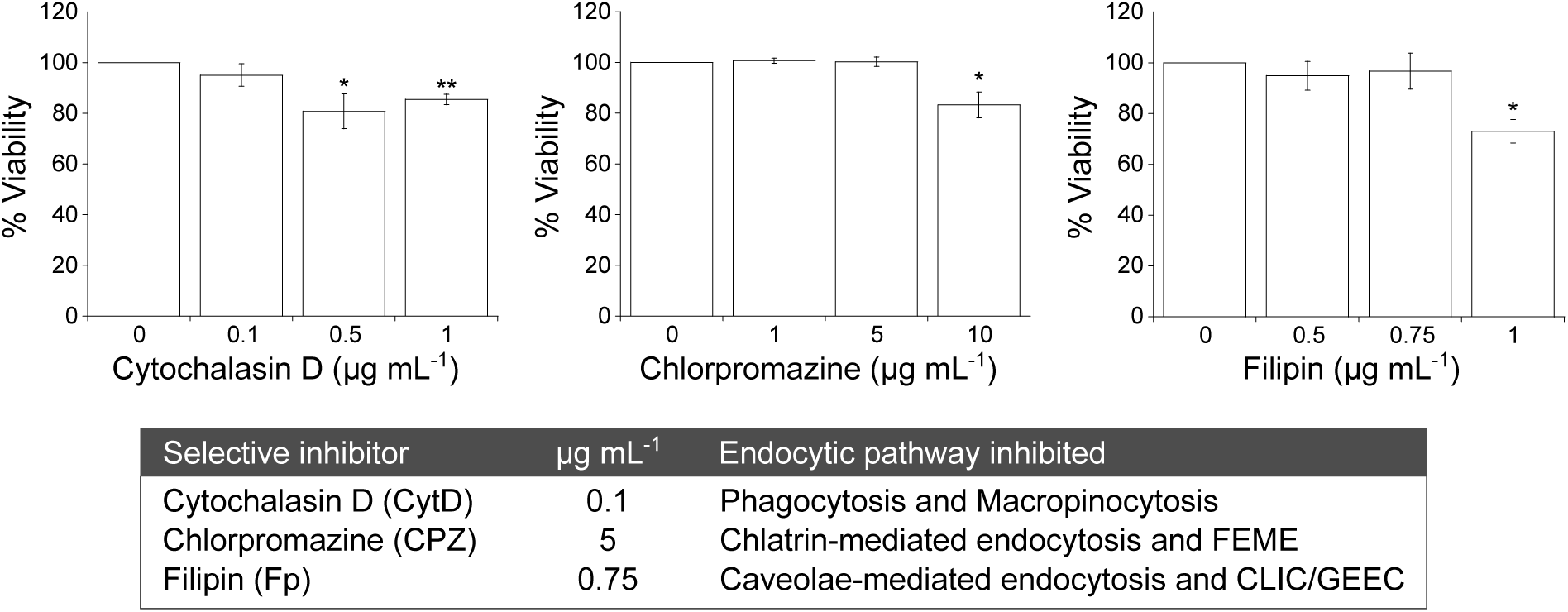
Viability of A549 cells after 2.5 hours in contact with increasing concentrations of endocytosis inhibitors. The percentages are normalized to control cells (without inhibitor). Data represent the mean ± standard deviation, n = 3. *, P < 0.05; **, P < 0.01. The concentrations of each inhibitor used for internalization experiments as well as the specific endocytosis routes inhibited by each compound are displayed on the table.

Then, the internalization pathways used by A549 cells to internalize MSR-1/magnetosomes were determined by using the aforementioned specific endocytosis inhibitors and flow cytometry. In addition, to inhibit all endocytic pathways, cells were co-incubated with MSR-1/magnetosomes at a sub-optimal temperature (4^◦^C), which reduces metabolic activity and thereby inhibits endocytosis, an energy-dependent process. The experimental setup is schematized in Figure 4. Briefly, A549 cells were cultured in four conditions: I) cells co-incubated with MSR-1/magnetosomes; II) cells co-incubated with MSR-1/magnetosomes at 4^◦^C; III) cells co-incubated with MSR-1/magnetosomes and with endocytosis inhibitors; and IV) control cells. After the incubation time, the cells corresponding to the four conditions were analyzed using flow cytometry. To be able to detect them, MSR-1 were labelled using rhodamine 123 that confers them with green fluorescence (λ_ex_ = 508 nm, λ_em_ = 528 nm). For the determination of magnetosome uptake by the cells, the side scattered light value was used, as nanoparticle-loaded cells have a higher value in side scattered light, probably as a result of the increased cellular complexity ^55^.

**Figure 4:**
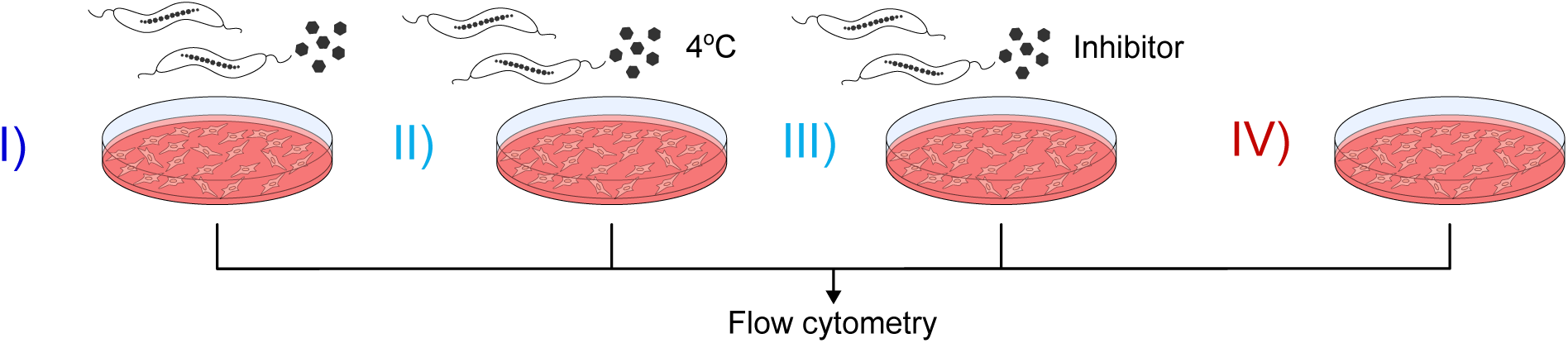
Scheme of the experiment performed to determine the internalization pathways used by A549 cells to uptake MSR-1/magnetosomes. Four sets of cells were cultured: I) MSR-1/magnetosome-loaded cells; II) MSR-1/magnetosome-loaded cells incubated at sub-optimal temperature (4 °C); III) MSR-1/magnetosome-loaded cells incubated with pharmacological inhibitors; and IV) control cells.

Figure 5 shows the histograms obtained by flow cytometry for the first of three replicates performed for each condition. The histograms of the three replicates are showcased on the Supplementary Information (Figures S3 and S4).

**Figure 5:**
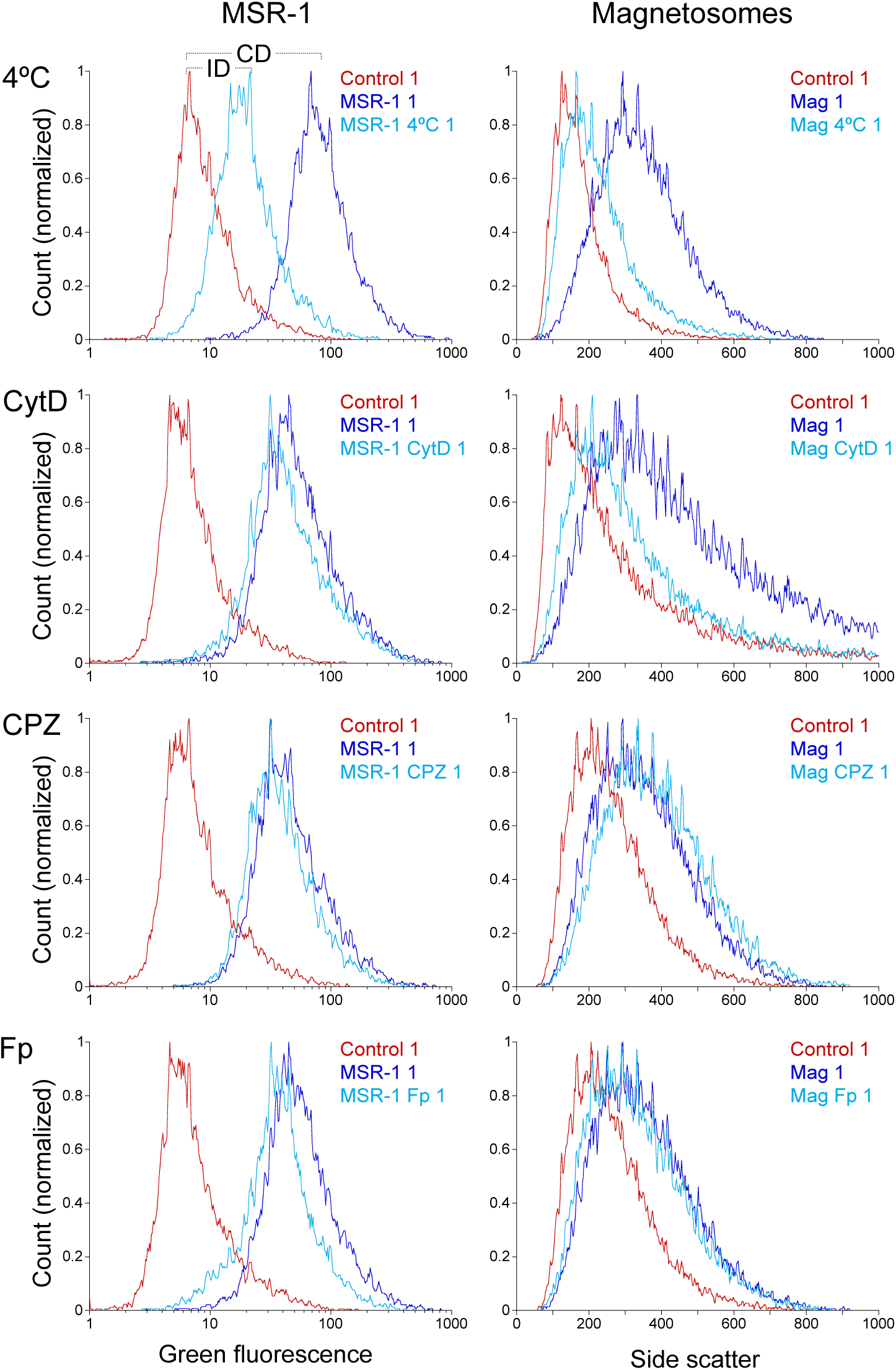
Examples of histograms showing measurements of A549 cells under the conditions listed in Figure 4A, obtained by flow cytometry. The mean values extracted from the histograms were used to calculate the inhibition percentage in each condition using Equation 1, allowing determination of the endocytic pathways employed by A549 cells to internalize MSR-1 (left) and magnetosomes (right). The differences in the mean values of green fluorescence/side scatter between loaded-cells and control cells (CD) as well as between loaded-cells + inhibitor and control cells (ID), are used in Equation 1 to calculate the percentage of endocytosis inhibition.

The mean values of the green fluorescence/side scatter of the cells in each condition were used to calculate the endocytosis inhibition percentages using Equation 1:

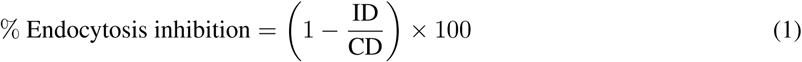

where CD (control difference) is the difference between the mean values of green fluorescence/side scatter of MSR-1/magnetosome-loaded cells (Figure 4I) and control cells (Figure 4IV) and ID (inhibition difference) is the difference between the mean values of green fluorescence/side scatter of MSR- 1/magnetosome-loaded cells in the presence of inhibitors (Figure 4II or Figure 4III) and control cells (Figure 4IV) (see example on Figure 5, MSR-1 at 4^◦^C) ^56^. The mean values used for the calculations as well as the endocytosis inhibition percentages can be found on the Supplementary Information (Tables S1 and S2). The endocytosis inhibition percentage results are represented on Figure 6. Here, it can be readily seen that the uptake of both MSR-1 and magnetosomes was significantly inhibited at 4^◦^C due to decreased metabolic activity, supporting endocytosis as the main mechanism of MSR-1/magnetosome internalization in A549 cells. The fact that the inhibition percentage did not reach 100% for both MSR-1 and magnetosomes is attributed to a small fraction (17% for MSR-1 and 24% for magnetosomes) remaining attached to the cell surface (as observed in Figure 1I for magnetosomes), which would be indistinguishable by flow cytometry.

**Figure 6:**
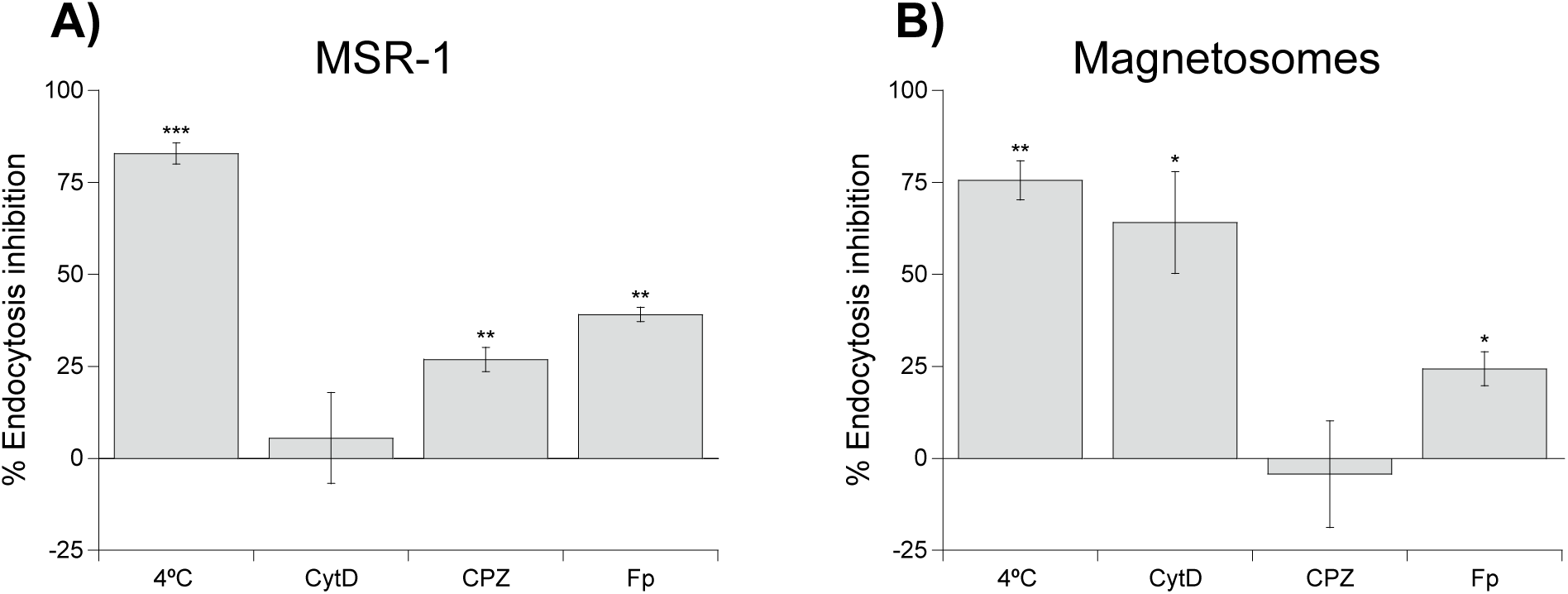
Inhibition of MSR-1/magnetosome endocytosis caused by sub-optimal temperatures or specific pharmacological inhibitors: cytochalasin D (CytD), chlorpromazine (CPZ) and filipin (Fp). Data represent the mean ± standard deviation, n = 3. *, P < 0.05; **, P < 0.01; ***, P < 0.001.

Regarding the endocytic pathways used by cells to internalize MSR-1, it might be intuitive to think that given their size (about 0.5 × 5 µm) the preferred endocytic pathway for cells to internalize them would be either phagocytosis or macropinocytosis as these pathways are involved in the uptake of large foreign bodies such as microorganisms ^48,49^. However, as shown in Figure 6A, cytochalasin D caused an almost negligible inhibition of MSR-1 internalization, while both chlorpromazine and filipin significantly inhibited MSR-1 internalization, proving this hypothesis wrong. From these results it can be inferred that A549 cells use receptor-mediated endocytosis to internalize MSR-1. It has been previously reported that certain proteins or molecules on the surface of bacteria such as *Listeria monocytogenes*, can actively influence host cell machinery to trigger a specific endocytic pathway in which is known as “the zipper mechanism”. More specifically, *Listeria* induces clathrin accumulation in the host cell surface followed by actin rearrangement and bacterial endocytosis ^58^. In view of the results, it could be argued that MSR- 1 has proteins or molecules on its surface that could cause the recruitment of specific proteins such as clathrin or caveolin to the host plasma membrane, subsequently inducing MSR-1 internalization into A549 cells by receptor-mediated endocytosis.

On the contrary, Figure 6B shows that magnetosome uptake is inhibited in the presence of cytochalasin D, suggesting that A549 cells can use phagocytosis or macropinocytosis to internalize magnetosomes. Although the nanoscopic size of isolated magnetosomes (around 40 nm) does not suggest this mechanism as the main one, magnetosomes tend to aggregate into clusters of around 1 µm, as previously shown in Figure 1I. Also in Figure 1I, it is observed that the endosomes containing magnetosome clusters are larger than 200 nm, which corresponds to the size of endosomes formed in phagocytosis and macropynocytosis ^48^. The fact that these two pathways do not depend on specific protein receptors may suggest that magnetosomes do not have as many specific ligands on their surface to target as MSR-1 do. In fact, they are intracellular structures that remain inside bacteria without any contact with the outer environment. Additionally, magnetosomes can also be internalized by caveolae-mediated endocytosis and/or CLIC/GEEC since there was a significant endocytosis inhibition when blocked by filipin. Clathrin-mediated endocytosis seems to not be involved in magnetosome internalization as the inhibition values obtained with this compound were null.

In a work by Wang et al. ^34^, where magnetosomes from *Magnetospirillum magneticum* AMB-1 were tested in HepG2 cells, a reduction of magnetosome uptake was reported when phagocytosis, macropinocytosis, and clathrin-mediated endocytic pathways were inhibited, but not when the caveolae-mediated endocytosis was blocked. One reason to explain the difference between Wang et al.’s work and the current study may be the use of different eukaryotic cells. Caveolae are present in many cell types, in some of which are of high density such as adipocytes, endothelial, and muscle cells, where they can account for more than 50% of the plasma membrane. However, there is a very low concentration of caveolae in some tissues such as the liver and absent in others such as the kidney proximal tubule ^60^. This could explain why the study from Wang et al. did not report any reduction in magnetosome uptake when caveolae-endocytic pathway was inhibited as the tested cells were from the liver, whereas the cells used for the present study are endothelial cells with higher density in caveolae.

## Conclusions

In this study, 3D imaging of *Magnetospirillum gryphiswaldense* (MSR-1) and isolated magnetosomes inside A549 cancer cells was performed for the first time using cryo soft X-ray tomography (cryo-SXT). This technique enabled direct visualization of MSR-1 and magnetosomes localized inside endosomes, without the need for complex sample preparation or staining, as required by other microscopy techniques previously used. Moreover, MSR-1 could be observed inside the cells in its entirety, not just through the presence of magnetosomes, as is often the case with TEM imaging.

Subsequently, by combining flow cytometry with specific endocytosis inhibitors, the endocytic pathways involved in the uptake of MSR-1 and magnetosomes were identified. The results show that MSR-1 primarily enter cells via receptor-mediated endocytosis, whereas magnetosomes are mainly internalized through receptor-independent mechanisms. This difference may be due to the presence of surface proteins on magnetotactic bacteria that promote specific interactions with the host cell’s endocytic machinery. By elucidating the uptake routes used by A549 cells, this work establishes a methodological framework for studying the internalization of magnetotactic bacteria and magnetosomes by target cells and advances their potential application in targeted cancer therapies.

## Materials and Methods

### Bacterial growth conditions and magnetosome isolation

*Magnetospirillum gryphiswaldense* strain MSR-1 (DSM 6361) was cultured at 28^◦^C without shaking for 48 hours in Flask Standard Medium (FSM) ^61^ containing (per litre of deionized water) 0.1 g KH_2_PO_4_, 0.15 g MgSO_4_×7 H_2_O, 2.38 g HEPES, 0.34 g NaNO_3_, 0.1 g yeast extract, 3 g peptone from soybean, 0.3% (wt/vol) of sodium pyruvate as the carbon source and 100 µM of Fe(III)-citrate. To stimulate magnetosome synthesis, MSR-1 were cultured in microaerobic conditions obtained by filling the culture bottles up to 3/4 and not closing them tightly. For the endocytosis experiments, 40 µM of rhodamine 123 (Sigma-Aldrich, R8004) was added to the FSM culture medium in order to make MSR-1 fluorescent.

Once collected by centrifugation (8,000 g, 15 minutes, 4^◦^C), the fluorescent bacteria were resuspended in PBS (1X, pH 7.4) and fixed with 2% glutaraldehyde overnight. Then, they were washed three times with PBS (1X, pH 7.4) before being resuspended in A549 culture media for the co-incubation experiments.

For magnetosome extraction and purification, bacteria were harvested by centrifugation (8,000 g, 15 minutes, 4^◦^C) after 48 hours of culture. The pellet was resuspended in 10 mL g^-1^ of 20 mM HEPES - 4 mM EDTA (pH 7.4) buffer. Bacteria were disrupted by French Press (GlenMills) applying 1,250 psig pressure. This step was repeated two times per sample to ensure cell disruption. Cell lysate was sonicated (40 W, 45 cycles of 15 s ON - 5 s OFF) at 4^◦^C to further segregate cell debris from magnetosomes. Magnetosomes were separated from other cell debris by a magnetic rack and resuspended in 10 mM HEPES - 200 mM NaCl (pH 7.4) buffer. Magnetosome suspension was further sonicated (40 W, 45 cycles of 15 s ON - 5 s OFF) at 4^◦^C and the magnetic rack washing was repeated. This step of sonicating and washing was performed three times. Finally, magnetosomes were collected in MilliQ water and stored at 4^◦^C.

### Eukaryotic cell culture

A549 human lung carcinoma cells ^62^ were cultured in RPMI-1640 medium (Sigma-Aldrich, R6504) supplemented with 10% fetal bovine serum (FBS) (Gibco, 10270-106), 2 mM L-glutamine (Sigma-Aldrich, 59202C) and a mixture of antibiotics (100 U ml^−1^ penicillin and 100 µg mL^−1^ streptomycin (Sigma-Aldrich, P4333)) and an antimycotic (0.25 µg mL^−1^ amphotericin B (Sigma-Aldrich, A2942)) at 37^◦^C in a humidified atmosphere (95% relative humidity) and 5% CO_2_.

### Cryo soft X-ray tomography (cryo-SXT)

Cryo transmission X-ray microscopy imaging was conducted at MISTRAL beamline in ALBA synchrotron (Barcelona, Spain) ^39,40^ under the awarded proposal 2022025668. A549 cells were grown in 6 well plates containing TEM grids (Quantifoil R2/2 holey carbon, gold) sterilized under UV (3 h) and functionalized with 0.01% poly-L-lysine (Sigma-Aldrich, P4707). After overnight incubation of the cells, 30 µg Fe_3_O_4_ mL^−1^ of magnetosomes or 5×10^9^ bacteria mL^−1^ were added in FBS-free RPMI medium and incubated for 2 h or 24 h, respectively. The grids were washed 3 times by immersion into PBS (1X, pH 7.4) before mounting them vertically into a Leica EM GP2 single-side blotting automated plunge freezer at 95% humidity. Once the grids were vertically mounted into the vitrobot, 2 µL of fiducials (BBI Group, concentrated 5X) were added onto the grid. The grids were then blotted from the back of the grid with filter paper for 3 seconds and quickly dropped into liquid ethane. Vitrified samples were kept under cryogenic conditions until they were measured in the MISTRAL beamline.

The samples were measured using a photon energy (520 eV) within the *water window* to take advantage of the natural absorption contrast of the biological material to acquire X-ray tomography data sets with the conditions described previously ^63^. The data sets were acquired using a zone plate objective with an outermost zone width of Δrn = 40 nm. The effective pixel size in the images was 13 nm. The image stacks were pre-processed to normalize and correct the intensity distribution delivered to the sample by the capillary condenser lens. Alignment and reconstruction of the tilted series was done by using IMOD software ^64^. The visualization and segmentation of the final volumes were carried out using Amira 3D and Chimera softwares.

### Flow cytometry: viability assay and MSR-1/magnetosome internalization experiments

Flow cytometry was used to establish the inhibitor concentrations that did not decrease cell viability. For this, cells were seeded in 6 well plates at a concentration of 5×10^5^ cells mL^-1^ and incubated overnight to attach. The next day, cells were incubated with increased concentrations of each inhibitor for 2.5 hours, washed once with PBS (1X, pH 7.4) and detached from culture plates by incubating them for 5 minutes with PBS 1X - 4mM EDTA. Cell viability was assessed by staining them with propidium iodide (Invitrogen, R37169), a red fluorescent stain that binds to DNA but can only penetrate the cells when their membranes are damaged, therefore staining only dead cells. Propidium iodide-stained cells were excited using a blue laser (488 nm) and recorded using the FL3 channel (620/20 nm). The inhibitors used in this study were cytochalasin D (Sigma-Aldrich, C8273), chlorpromazine (Sigma-Aldrich, 31679), and filipin (Sigma-Aldrich, F9765).

For the MSR-1/magnetosome internalization experiments, cells were seeded at a concentration of 5×10^5^ cells mL^-1^ in 6 well plates and incubated overnight. The next day, the cells were pre-incubated for 30 minutes either at 4^◦^C or with the corresponding inhibitors for 30 minutes and subsequently put in contact with fluorescent MSR-1 (5×10^9^ bacteria mL^-1^) or magnetosomes (30 µg Fe_3_O_4_ mL^-1^) in FBS-free culture medium for 2 hours. Then, the culture medium was removed and the cells were washed once with PBS and detached from the culture plates by incubating them for 5 minutes with PBS - 4mM EDTA in the cell incubator in order to perform flow cytometry measurements. For MSR-1 internalization experiments, as the bacteria were fluorescent due to the rhodamine 123, the samples were excited using the blue laser of the cytometer (488 nm) and the fluorescence was recovered in the FL1 channel corresponding to green fluorescence (525/40 nm). For magnetosome internalization, the side scattered light was the indicator of the internalization as explained on the main text.

The equipment used was a Beckman Coulter Gallios cytometer of the Analytic and High Resolution Microscopy in Biomedicine Service (SGIker) of the University of the Basque Country (UPV/EHU).

## Acknowledgements

This work was supported by the grant PID2023-146448OB-C21 funded by MICIU/ AEI / 10.13039/501100011033 / FEDER, UE and by the grant IT1479-22 funded by the Basque Government. L.G. would like to thank the financial support from the grant PRE2018-083255 funded by MCIN/AEI/ 10.13039/501100011033 and by European Union NextGenerationEU/PRTR and from the European Union’s Horizon Europe research and innovation programme under the Marie Skłodowska-Curie grant agreement No. 101150206 (DroneMTB). The authors acknowledge awarded ALBA synchrotron beamtime (proposal 2022025668), and the BL09-MISTRAL beamline staff for assistance in cryo-SXT experiments. The authors would like to extend their gratitude to Dr. Adriana Rojas and Idoia Iturrioz from CIC bioGUNE for their technical support in sample preparation for cryo-SXT measurements, as well as to Dr. Eva Pereiro and Dr. Daniel Chevrier for their valuable guidance and support in preparing and conducting the experiments.

