## Supplementary Information for "Identifying the internalization pathways of magnetotactic bacteria and magnetosomes by cancer cells"

[5] Dpto. Electricidad y Electrónica, Universidad del País Vasco - UPV/EHU, 48940 Leioa, Spain

[6] Dpto. Física Aplicada, Universidad del País Vasco - UPV/EHU, 48013 Bilbao, Spain

November 15, 2025

### Transmission electron microscopy images of MSR-1 and magnetosomes

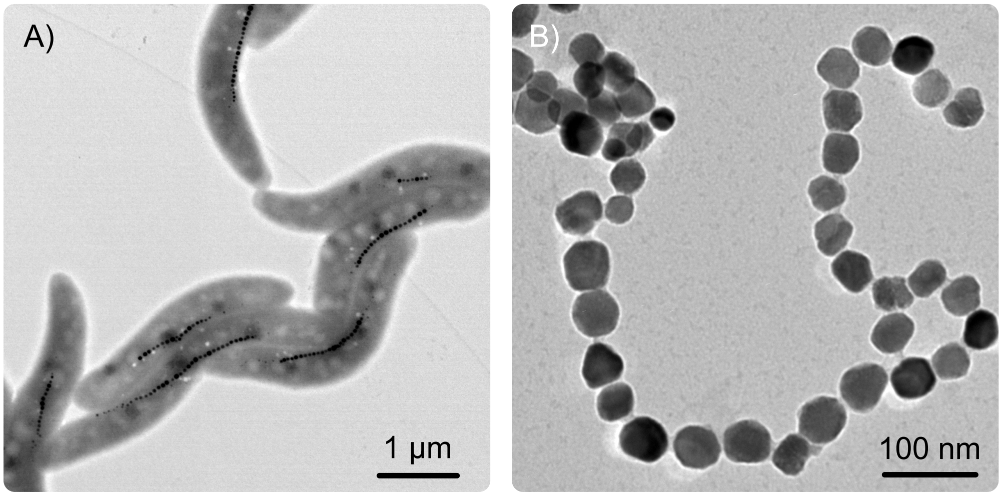


Figure S1: Transmission electron microscopy images of *Magnetospirillum gryphiswaldense* MSR-1 bacteria (A) and isolated magnetosomes (B). The images were acquired with a JEOL JEM-1400 Plus electron microscope at an accelerating voltage of 120 kV in the Analytic and High Resolution Microscopy in Biomedicine Service (SGIker) of the University of the Basque Country (UPV/EHU).

### Cryo soft X-ray tomography images and reconstructions of A549 cells containing MSR-1 and magnetosomes

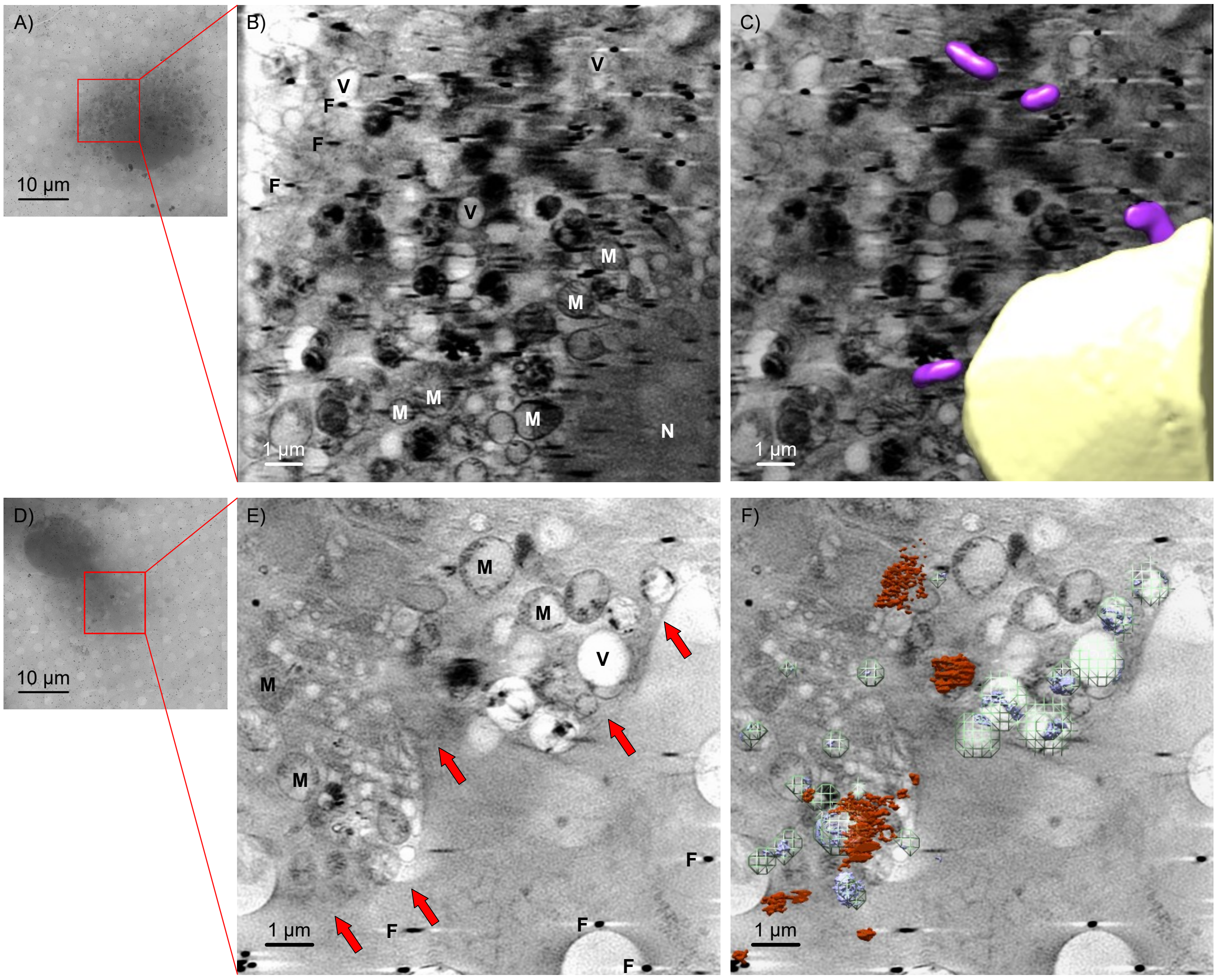


Figure S2: Cryo-SXT images of A549 cells loaded with MSR-1 (A-C) and magnetosomes (D-F). A,D) X-ray images of the measured cells, with the region chosen for tomogram acquisition highlighted by a red square. B,C,E,F) Cryo-SXT tomogram slices (B,E) and volumetric representation (C,F) of the tomogram highlighting cell nuclei (yellow), MSR-1 (magenta, C), magnetosomes inside endosomes (blue, F), endosomes (green mesh, F), and magnetosomes outside or entering the cell (red, F). The observed intracellular structures are marked as: N, cell nuclei; M, mitochondria; V, vesicles; F, fiducials; red arrows, plasma membrane. The videos of the volumetric reconstructions are shown in Supplementary Videos as S3 and S4.

### Histograms obtained by flow cytometry as a result of the endocytosis inhibition experiments

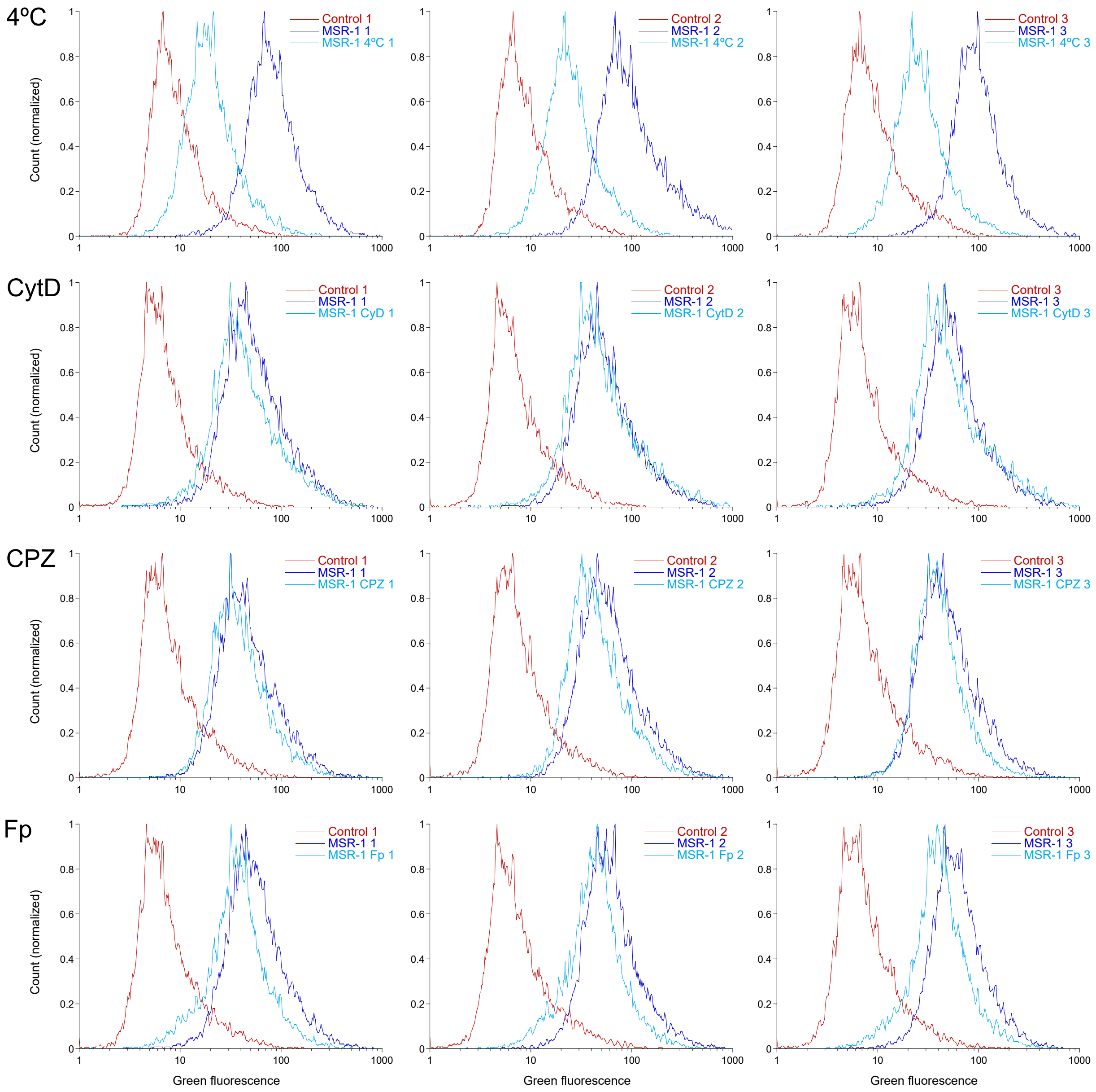


Figure S3: Histograms showing the green fluorescent values of control A549 cells (Figure 4IV), A549 cells incubated with MSR-1 (Figure 4I), and A549 cells incubated with endocytosis inhibitors and MSR-1 (Figure 4II,III). The inhibitors used are sub-optimal temperature (4°C), cytochalasin D (CytD), chlorpromazine (CPZ), and filipin (Fp). The mean values of the green fluorescence used to calculate the MSR-1 inhibition percentages are shown in Table S1.

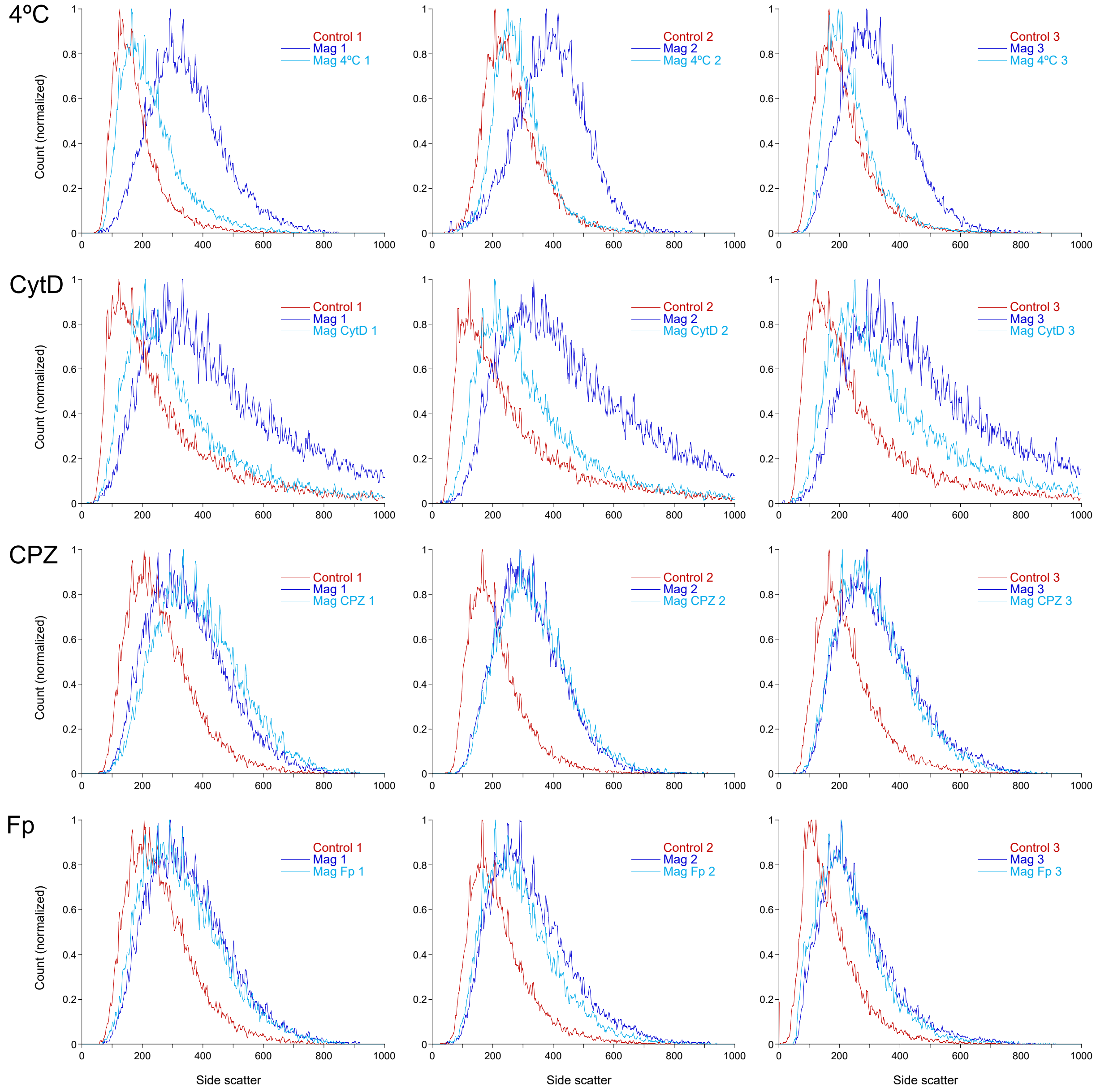


Figure S4: Histograms showing the side scattered light values of control A549 cells (Figure 4IV), A549 cells incubated with magnetosomes (Figure 4I), and A549 cells incubated with endocytosis inhibitors and magnetosomes (Figure 4II,III). The inhibitors used are sub-optimal temperature (4°C), cytochalasin D (CytD), chlorpromazine (CPZ), and filipin (Fp). The mean values of the side scatter used to calculate the magnetosome inhibition percentages are shown in Table S2.

| MSR-1 internalization | | | | | |
|---|---|---|---|---|---|
| Inhibitor | Replicate | Control | MSR-1 | MSR-1 + inhibitor | % Inhibition |
| 4°C | 1 | 11.56 | 99.57 | 25.25 | 84.44 |
| | 2 | 11.32 | 142.69 | 31.55 | 84.60 |
| | 3 | 12.16 | 115.68 | 33.24 | 79.64 |
| | | | | Mean | 82.89 |
| | | | | SD | 2.82 |
| CytD | 1 | 9.43 | 71.56 | 59.30 | 19.73 |
| | 2 | 10.44 | 72.98 | 74.13 | -1.84 |
| | 3 | 10.58 | 75.62 | 76.55 | -1.43 |
| | | | | Mean | 5.49 |
| | | | | SD | 12.34 |
| CPZ | 1 | 10.70 | 60.38 | 48.94 | 23.03 |
| | 2 | 10.19 | 77.12 | 57.99 | 28.58 |
| | 3 | 11.07 | 63.66 | 48.47 | 28.88 |
| | | | | Mean | 26.83 |
| | | | | SD | 3.30 |
| Fp | 1 | 10.10 | 72.06 | 46.60 | 41.09 |
| | 2 | 10.76 | 79.42 | 53.85 | 37.24 |
| | 3 | 10.38 | 77.43 | 51.40 | 38.82 |
| | | | | Mean | 39.05 |
| | | | | SD | 1.94 |

Table S1: Green fluorescence mean values obtained from the histograms in Figure S3 used in Equation 1 to calculate the inhibition percentages highlighted in orange and represented in Figure 6.

| Magnetosome internalization | | | | | |
| --- | --- | --- | --- | --- | --- |
| Inhibitor | Replicate | Control | Magnetosomes | Magnetosomes + inhibitor | % Inhibition |
| 4°C | 1 | 185.38 | 355.41 | 236.42 | 69.98 |
| | 2 | 272.10 | 404.60 | 297.97 | 80.48 |
| | 3 | 220.17 | 332.27 | 246.78 | 76.26 |
| | | | | Mean | 75.57 |
| | | | | SD | 5.28 |
| CytD | 1 | 311.38 | 513.41 | 367.75 | 72.10 |
| | 2 | 310.02 | 529.58 | 371.61 | 71.95 |
| | 3 | 314.33 | 546.76 | 434.73 | 48.20 |
| | | | | Mean | 64.08 |
| | | | | SD | 13.75 |
| CPZ | 1 | 177.62 | 260.75 | 272.32 | -13.92 |
| | 2 | 220.95 | 332.74 | 345.33 | -11.26 |
| | 3 | 235.60 | 338.60 | 325.98 | 12.25 |
| | | | | Mean | -4.31 |
| | | | | SD | 14.40 |
| Fp | 1 | 270.77 | 362.40 | 342.78 | 21.41 |
| | 2 | 229.90 | 334.99 | 303.79 | 29.69 |
| | 3 | 177.95 | 263.86 | 244.94 | 22.02 |
| | | | | Mean | 24.37 |
| | | | | SD | 4.61 |

Table S2: Side scatter mean values obtained from the histograms in Figure S4 used in Equation 1 to calculate the inhibition percentages highlighted in orange and represented in Figure 6.
